# Challenges and costs of asexuality: Variation in premeiotic genome duplication in gynogenetic hybrids from *Cobitis taenia* complex

**DOI:** 10.1101/2021.07.15.452483

**Authors:** D. Dedukh, A. Marta, K. Janko

## Abstract

The transition from sexual reproduction to asexuality is often triggered by hybridization. The gametogenesis of many hybrid asexuals involves a stage of premeiotic genomic endoreduplication leading to the production of clonal gametes and bypassing genomic incompatibilities that would normally cause hybrid sterility. However, it is still not clear at what gametogenic stage the endoreplication occurs, how many gonial cells it affects and whether its rate differs among clonal lineages. Here, we investigated meiotic and premeiotic cells of diploid and triploid hybrids of spined loaches (Cypriniformes: *Cobitis*) that reproduce by gynogenesis. We found that naturally as well as experimentally produced F1 hybrid strains undergo an obligatory genome duplication event to achieve asexuality, occurring in the gonocytes just before entering meiosis or, rarely, one or few divisions before meiosis. Surprisingly however, the genome endoreplication was observed only in a minor fraction of the hybrid’s gonocytes, while the vast majority were unable to duplicate their genomes and consequently could not proceed beyond pachytene due to defects in pairing and bivalent formation. We also noted that the rate of endoreplication was significantly higher among gonocytes of hybrids from successful natural clones than of experimentally produced F1 hybrids, indicating that interclonal selection may favour lineages which maximize the rate of premeiotic endoreduplication. We conclude that asexuality and hybrid sterility are intimately related phenomena and the transition from sexual reproduction to asexuality must overcome significant problems with genome incompatibilities with possible impact on reproductive potential.

## Introduction

Species are fundamental evolutionary units, presumably evolving in a continuum from intermixing populations to independent entities isolated from other species by pre- and postzygotic barriers (Avise, 2005; Coyne et al., 2004). Their formation is thus frequently accompanied by interspecific hybridization which may have positive (Abbott et al., 2013; Coyne et al., 2004; Mallet, 2005; Rieseberg and Willis, 2007), as well as negative impacts (Arnold and Hodges, 1995; Coyne et al., 2004; Rieseberg, 2001) and appears to be a mighty evolutionary force. With incompatibilities accumulating among their genomes, the crossing of parental species often affects the fertility of hybrids ultimately leading to their sterility (Coyne and Orr, 1998; Maheshwari and Barbash, 2011). Hybrid sterility may have various causes (Geraldes et al., 2006; Payseur et al., 2004) and its molecular underpinning is still little understood. However, an important cause of hybrid sterility is the improper pairing and recombination between orthologous chromosomes from two different parental species during meiotic prophase, leading to the abruption of meiosis and/or aneuploid gametes (Arnold and Hodges, 1995; Coyne et al., 2004; Rieseberg, 2001). Nonetheless, hybridisation not only affects interactions between the two admixed genomes but it also can modify gametogenic pathways and induce the switch of a hybrid’s reproduction towards asexuality (Abbott et al., 2013; Bullini, 1994; Choleva et al., 2012; Ernst, 1918; Lenormand et al., 2016; Stenberg and Saura, 2013).

Traditionally, sex and asexuality have been viewed as contrasting dichotomies, but in reality, they rather represent two extremes of a continuum. Indeed, even sexual organisms can sometimes spontaneously produce unreduced gametes (Brownfield and Köhler, 2011; Lampert, 2008; Mason and Pires, 2015) and asexual organisms in fact represent a very diverse group that employ a wide spectrum of cytological mechanisms for gamete production. These range from completely ameiotic processes (apomixis) to those involving more or less distorted meiotic divisions (automixis) (Neaves and Baumann, 2011; Stenberg and Saura, 2013, 2009). Surprisingly, although sexual and asexual reproduction represent a major and intensively studied paradox of evolutionary biology (Avise, 2021, 2005; Otto and Lenormand, 2002), very little is known about the cellular and molecular machinery causing the alterations between both reproduction types. Detailed investigation of these pathways could give an answer to basic questions such as what mechanisms cause the transitions from sexual reproduction to asexuality? Why some types of gametogenic aberrations are more common than others? And, what challenges do asexuals face during the alterations of cellular pathways?

At least in hybrid asexuals, it appears that the switch from sexual to clonal reproduction involves the modification of conservative gametogenic pathways, which is already occurring in the F1 generation (Lenormand et al., 2016; Neaves and Baumann, 2011; Stenberg and Saura, 2013, 2009). Interestingly, in cases where hybridization leads to aberrant chromosomal pairing and hybrid sterility, the adoption of clonal gametogenesis may partially overcome such conflicts and restore fertility (Figure 1) (Dedukh et al., 2020; Janko et al., 2018; Neaves and Baumann, 2011; Stenberg and Saura, 2013). One example of this is premeiotic endoreplication, which is a widespread gametogenic alteration found among a vast variety of unrelated asexual organisms such as plants, invertebrates and vertebrates (Figure 1) (Stenberg and Saura, 2009; Storme and Geelen, 2013; Suomalainen, 1987). During this process chromosomes of gonial cells are duplicated ensuring successful progression through meiosis due to bivalents forming between identical copies of chromosomes (Dedukh et al., 2015; Dedukh et al., 2020; Kuroda et al., 2018; Lutes et al., 2010; Macgregor and Uzzell, 1964). As a result, endoreplication rescues the hybrid’s fertility and ensures clonal propagation of the maternal genome (Figure 1) (Dedukh et al., 2020; Kuroda et al., 2019). Hybrid asexuality and sterility thus share some common cytological ground; both tend to emerge in hybrids between substantially diverged species rather than between closely related ones (Bateson, 1909; Ernst, 1918; Janko et al., 2018; Moritz, 1989; Russell, 2003).

**Fig 1.**
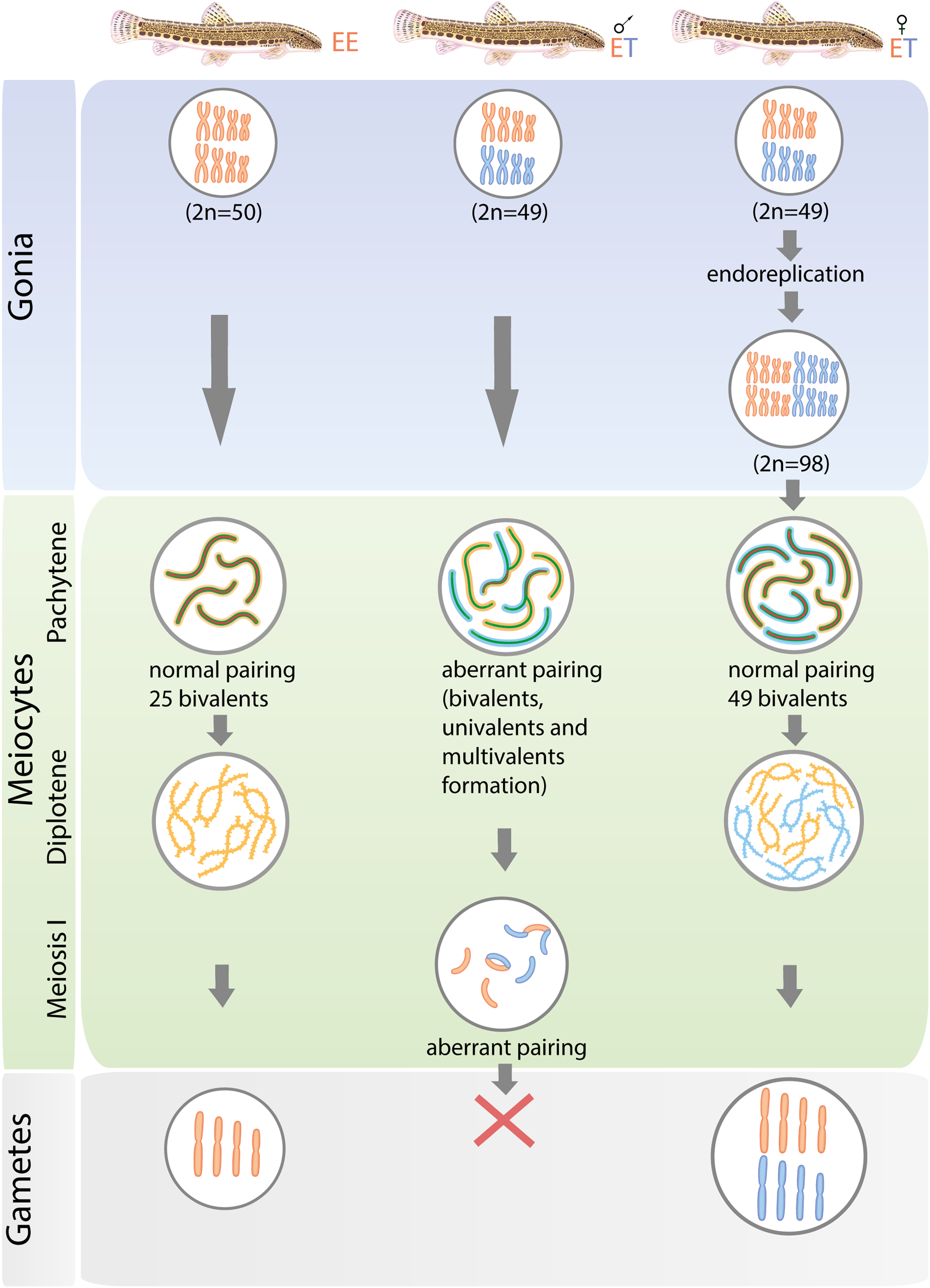
Schematic overview of gametogenic pathway during sexual reproduction (a), hybrid sterility (b) and asexuality (c). Each column shows individuals, gametogenic pathways with the indication of the germ cells, cells in two meiotic stages (pachytene and diplotene) and gametes. A (orange chromosomes), B (blue chromosomes) indicates genomes of different parental species. (a) During sexual reproduction, gonia cells enter meiosis, which results in a haploid egg. Homologous chromosomes are joined by the synaptonemal complex (green color indicates the lateral element of synaptonemal complexes; red color indicates the central element of synaptonemal complexes) during pachytene which is disassembled by the diplotene stage of meiosis. (b) Hybrid sterility caused by aberrant chromosome pairing during pachytene (modified from Dedukh et al., 2020). (c) Reproduction of asexual hybrids is realized by endoreplication of the genomes in gonial cells allowing bivalent formation between identical chromosomal copies resulted in diploid gamete formation (modified from Dedukh et al., 2020).

In-depth studies of asexual gametogenesis are relatively rare but it has been hypothesized that once a successful clone emerges most of its germ cells follow the same pathway towards production of unreduced gametes (Moritz, 1989). Surprisingly though, studies on laboratory synthesized hybrids between medaka fish species as well as on parthenogenetic lizards from genus *Aspidoscelis* both showed that endoreplication occurred only in a small portion of hybrid’s germ cells, while most of their oogonia failed to endoreplicate their genome and did not develop to gametes (Hamaguchi and Sakaizumi, 1992; Newton et al., 2016; Shimizu et al., 2000).

In this study, we aim to analyse the gametogenesis of natural clonal lineages as well as laboratory-induced hybrids in order to test whether premeiotic genome endoreplication is indeed occurring in all or the majority of the hybrid’s germ cells and if even successful natural clones face problems in gamete production inherent to their hybrid origin.

We focused on spined loaches *Cobitis* (Cypriniformes: Cobitidae) which are an excellent model to investigate the emergence and evolutionary consequences of asexuality. Three species of these freshwater fishes meet and reproductively interact in Europe: *C. taenia* with diploid karyotype involving 2n=48 chromosomes in somatic tissues (henceforth its genome will be ‘T’, so that diploid pure species is denoted ‘TT’), *C. elongatoides* (EE, 2n=50) and *C. tanaitica* (NN, 2n=50) (Bohlen and Ráb, 2001; Janko et al., 2007; Majtánová et al., 2016; Marta et al., 2020). Previous studies showed that their hybridization produces hybrids both in laboratory and natural conditions (Choleva et al., 2012; Janko et al., 2018). Male hybrids between *C. elongatoides* and *C. taenia* are sterile due to the aberrant pairing of their chromosomes (Figure 1) (Dedukh et al., 2020; Juchno and Boroń, 2006). Regardless of this, the fertility of the hybrid females is sustained by premeiotic genome duplication and consequently, diploid females (of ET or EN genetic constitution with 49 and 50 chromosomes, respectively) reproduce gynogenetically by clonal eggs (Figure 1) (Dedukh et al., 2020; Juchno et al., 2016). Occasionally their eggs incorporate sperm from parental species leading to the establishment of triploid *C. elongatoides-taenia* (ETT, 3n=73) and *C. elongatoides-tanatitica* (EEN, 3n=75) gynogenetic females (Bohlen and Ráb, 2001; Dedukh et al., 2020; Majtánová et al., 2016; Marta et al., 2020). The hybridization among these species has been dynamic since the Pleistocene and has led to high clonal diversity. Many clones originated relatively recently during the Holocene and occur in secondary hybrid zones between *C. elongatoides* and *C. taenia* or *C. tanaitica*, but one successful clonal lineage, EEN, colonised vast areas over Europe since its ancient origin approximately 300 kya (Janko et al., 2012, 2005).

Here, we analysed the gametogenic pathways of both experimental F1 and naturally occurring clonal linages and tested the proportion of their cells which successfully develop into clonal gametes. Specifically, we focused on oocytes during the pachytene and diplotene as well as gonocytes and analysed their distribution throughout ovaria of asexual hybrid biotypes, including diploid, triploid, newly synthesized F1’s as well as successfully established natural clonal lineages. We also estimated the ploidy of individual germ and meiotic cells using species polymorphic and chromosome-specific FISH probes. In each type of hybrid, we investigated how its cells passed through meiotic checkpoints and checked the ability of their germ cells to undergo premeiotic duplication of the genomes leading to viable gametes.

## Results

### All diplotene oocytes have a duplicated genome and properly paired chromosomes with bivalents

Somatic cells of diploid ET hybrids have 2n=49 chromosomes, while triploid hybrids with ETT and EEN genome composition have 3n=73 and 75 chromosomes, respectively (Bohlen and Ráb, 2001; Majtánová et al., 2016; Marta et al., 2020). Consequently, after endoreplication, their germ cells should have 98, 146 and 150 chromosomes respectively, leading to 49, 73 and 75 bivalents in oocytes of diploid ET and triploid ETT and EEN hybrids (this study; Dedukh et al., 2020). To test if the germ cells of diploid (natural and laboratory generated F1 hybrids) as well as triploid hybrids showed evidence of genome duplication, we determined the number of bivalents in oocytes during the diplotene stage.

From a total of 263 observed oocytes, all possessed the expected number of bivalents / chromosomes (Figure 2, 3a; Supplementary Table 1; Supplementary Figure S1a, S2a), indicating that a hybrid’s oocytes at the diplotene stage possess a duplicated number of chromosomes compared to somatic cells. We did not observe any sign of abnormal pairing or the presence of multivalents and univalents. To discern whether the bivalents formed by pairing of homoeologous (ExE and TxT, or NxN) or orthologous (ExT or ExN) chromosomes, we identified bivalents by morphology using previously constructed lampbrush chromosome maps (Dedukh et al., 2020). As a result, we noticed that all bivalents where the aforementioned method was applicable, strictly corresponded to homoeologues, confirming the results of Dedukh et al. (2020). In addition, we also used FISH identification with species polymorphic satellite markers (satCE02), which is known to occur on two chromosomes in *C. taenia* and a single chromosome in *C. elongatoides* (Supplementary Figure 3c,d,g,h) (Marta et al., 2020). Using the satCE02 marker, in diploid hybrids we detected 2 bivalents of *C. taenia* and 1 bivalent of *C. elongatoides* while in triploid hybrids we identified 4 bivalents of *C. taenia* and 1 bivalent of *C. elongatoides* (Figure 3b-f; Supplementary Figure 1b-d). These observations indicate that diplotenic oocytes have their chromosomal sets premeiotically duplicated and that pairing occurs between homoeologous chromosomes (Figure 3g; Supplementary Figure 1e, 2b).

**Fig 2.**
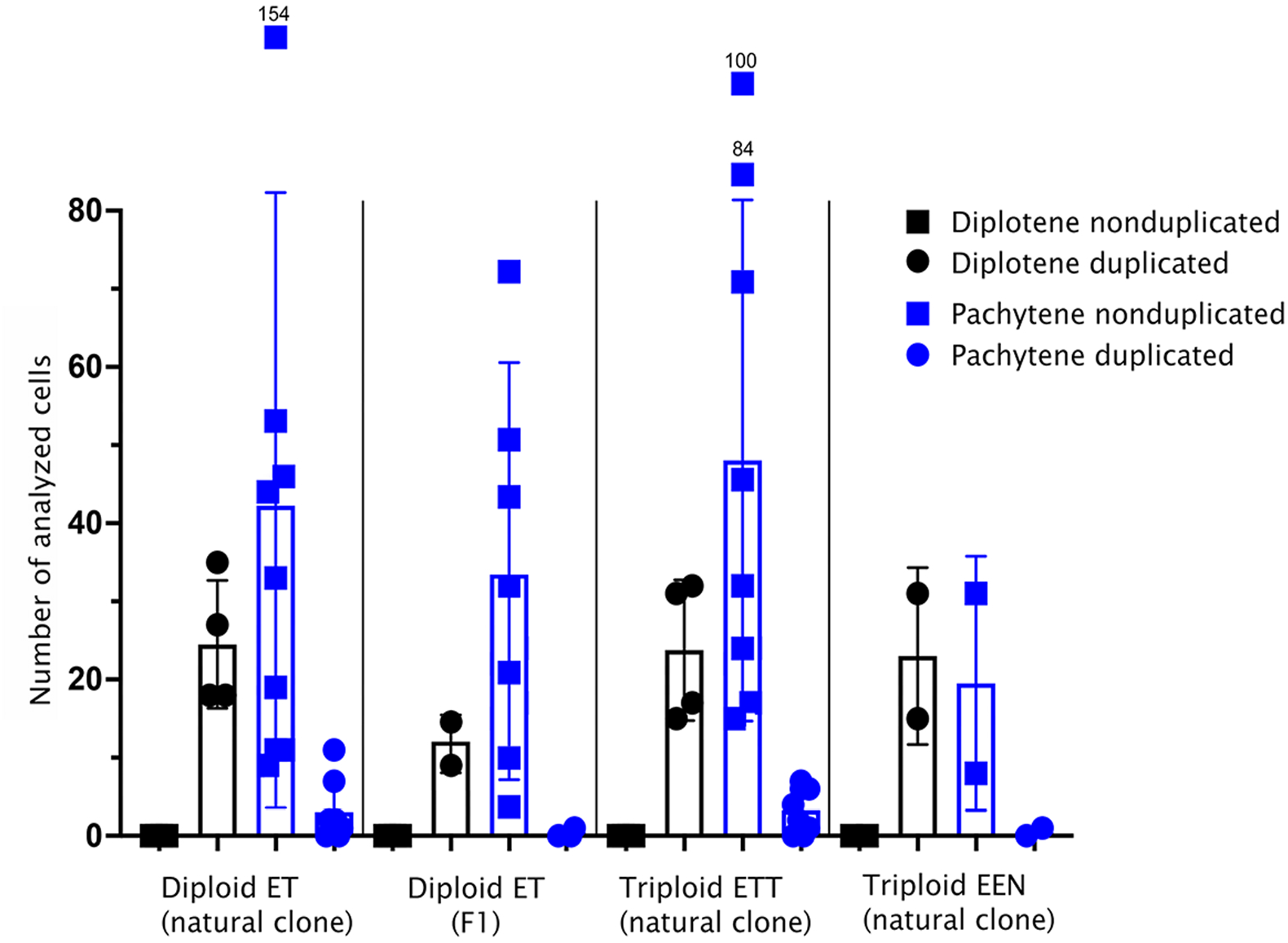
The relative number of cells with duplicated and notduplicated genomes observed during pachytene and diplotene in natural diploid ET hybrids, artificial F1 diploid hybrids, and natural triploid ETT and EEN hybrids.

**Fig 3.**
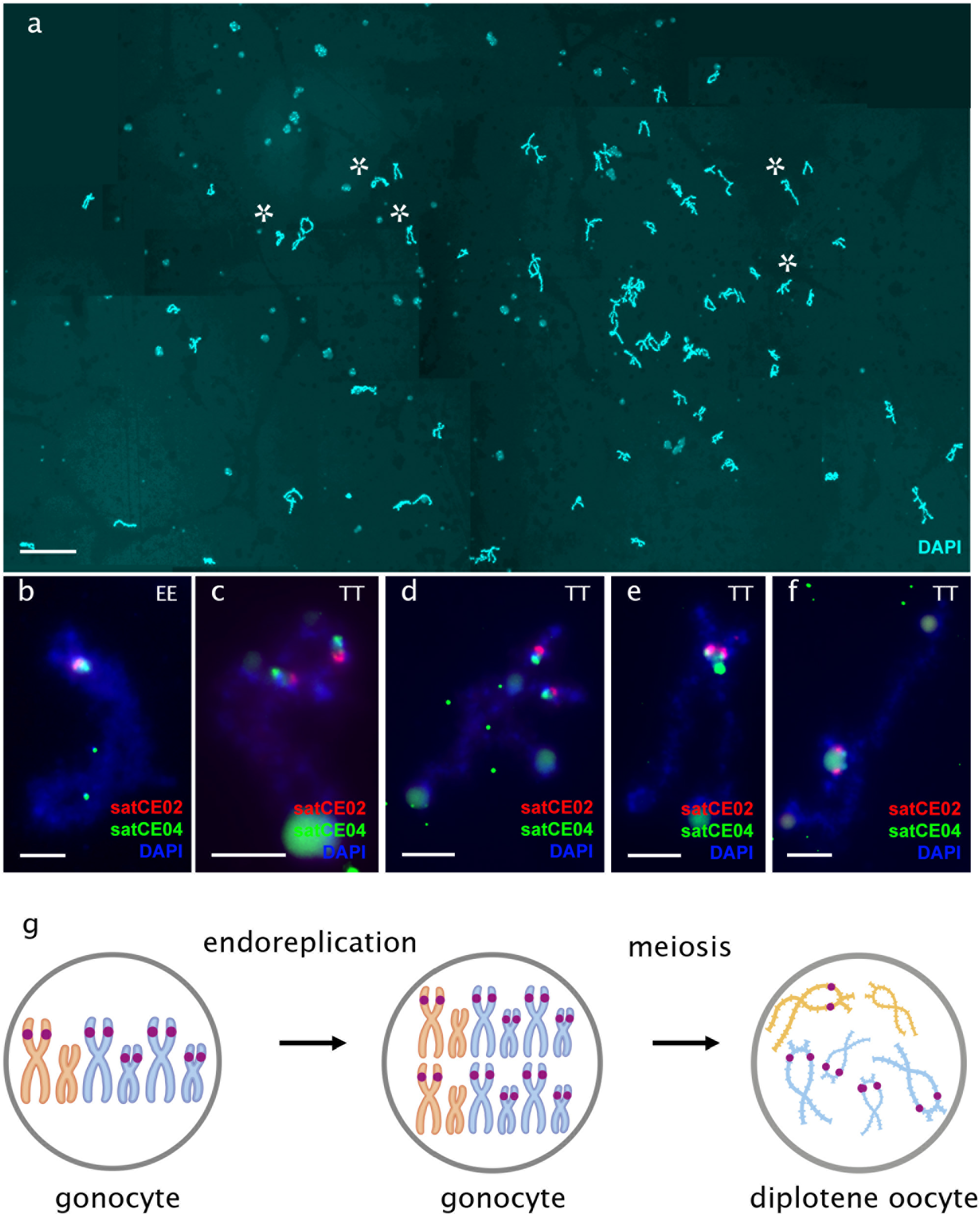
FISH based identification of bivalents from diplotene oocyte of triploid ETT female. (a) Full diplotene chromosomal spread from the individual oocyte including 73 bivalents. Since the chromosomal spread from individual oocyte was large, five images were taken and merged into one. Asterisks indicate enlarged bivalents represented on b-f panel. Scale bar = 50 μm. (b-f) High-resolution mapping of species polymorphic (satCE02, red) and centromeric markers (satCE4, green) on diplotene chromosomes (so-called “lampbrush chromosomes”) indicates one bivalent of *C. elongatoides* (b) and 4 bivalents of *C. taenia* (c-f). Scale bar = 50 μm. (g) Schematic representation of gametogenic pathway, which results in the formation of bivalents observed during diplotene. Purple marks indicate bivalents identified by FISH with species polymorphic marker satCE02 as well as presumptive karyotype composition in gonocytes.

### Pachytene fall into two types: those with duplicated chromosomes and proper bivalents and those with few bivalents and many univalents

In contrast to diplotenic cells, we found a surprising variability in the number of paired homologs in oocytes of the pachytene stage. Using immunofluorescence detection of central (SYCP1) and lateral (SYCP3) components of synaptonemal complexes (Saito et al. 2014; Blokhina et al. 2019) we observed that in triploid ETT hybrids, only ~8% (n = 26) of pachytene chromosome spreads had 73 bivalents as would be expected after premeiotic endoreduploication (Figure 2, 4b, Supplementary Table 1). The vast majority of their cells possessed only 20-23 bivalents and several unpaired or partially paired chromosomes (Figure 2, 4a, Supplementary Table 1). In a widespread triploid EEN clone, only one pachytene cell in one individual out of 40 scored spreads (~1.5%) had duplicated sets of chromosomes (Figure 2, Supplementary Table 1; Supplementary Figure S4a,b).

**Fig 4.**
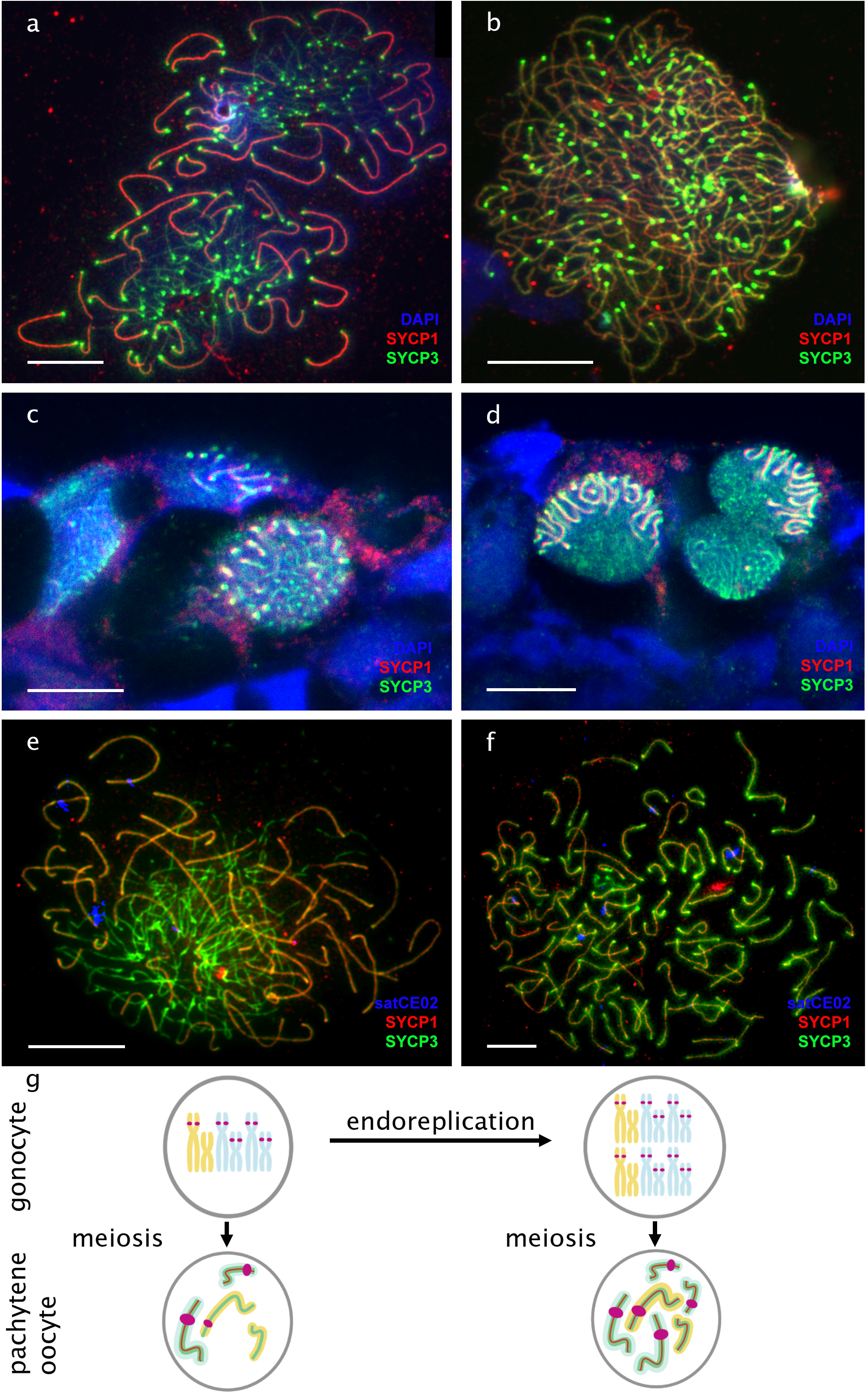
Meiotic cells at pachytene stage from ovaries of triploid ETT females. Visualization of synaptonemal complexes using immunolabeling with antibodies against SYCP3 protein (green) and SYCP1 (red) stained with DAPI (blue) on pachytene chromosomal spreads (a,b) and whole tissues (c,d). Images c-d are single confocal sections of 0.6 μm in thickness. Identification of bivalents (accumulating both SYCP3 and SYCP1) and univalents (accumulating only SYCP3) on pachytene spreads exhibiting the presence of two population of cells differed in ploidy level: triploid, with both bivalents and univalents (a, c) and hexaploid, only with bivalents (b, d). (e, f) Mapping of species polymorphic marker (satCE02, blue) on cells with both bivalents and univalents (e) as well as bivalents only (f). (g,h) Schematic representation of gametogenic pathways which result in bivalent formation in two populations of pachytene cells. Purple marks indicates bivalents identified by FISH with species polymorphic marker satCE02 as well as presumptive karyotype composition in gonocytes. Scale bar = 10μm.

A similar pattern was seen in natural ET diploids, with ~6% pachytene spreads containing 49 bivalents (n = 24) and the vast majority of cells showing a combination of 12-15 bivalents, 1-2 multivalents and several univalents (Figure 2; Supplementary Table 1; Supplementary Figure 5a,b). The incidence of cells with proper bivalents during pachytene was even lower in experimental F1 hybrids where only one cell out of 299 showed a signal of 49 properly formed bivalents after endoreplication (Figure 2; Supplementary Table 1).

The differences between hybrid biotypes (F1 ET, natural ET, natural ETT and natural EEN) in the proportion of duplicated and nonduplicated pachytene cells was highly significant (generalized linear model with binomial error distribution, p.val =13*10^-5). Post hoc tests further showed significant differences among pairs of biotypes containing the comparisons between F1 ET and natural ET (p.val =9*10^-5) or ETT (p.val =39*10^-5). On the other hand, no significant differences were observed between pairs containing F1 ET and natural EEN, neither between EEN and natural ET or ETT, nor between natural ET and ETT. In summary, we may conclude that F1 ET hybrids had a significantly lower proportion of duplicated cells as compared to their natural counterparts, while in EEN the low number of scored cells prevented any clear-cut conclusion to be made.

To verify previous observations on pachytene spreads, we also performed whole mount immunofluorescent staining on pachytene meiocytes inside entire gonadal fragments. We did so by investigating the gonads of diploid and triploid hybrid females using the same antibodies against SYCP1 and SYCP3, and similar to previous analyses, we also detected two types of pachytene cells; those with only bivalents and those containing a mixture of univalents and bivalents (Figure 2; Figure 4c, d; Supplementary Table 1). Interestingly, the analysis of entire gonads indicated that pachytene cells containing only bivalents were not organized in clusters but were surrounded by cells with both bivalents and univalent (Figure 4c, d).

### Pachytene oocytes containing bivalents and univalents do not have a duplicated genome, while those with only bivalents do

To discern whether the two cell types in pachytene differ in number of genomic copies, we determined their ploidy by applying FISH with the species polymorphic satellite marker satCE02 to pachytene chromosomal spreads (Marta et al., 2020). We observed that triploid’s pachytene oocytes with exclusively bivalents contained satCE02 signals on 5 bivalents; 4 corresponding to *C. taenia* bivalents and the other corresponding to the *C. elongatoides* bivalent (Figure 4f). These results matched our observations of diplotene using the same FISH marker (see above; Figure 3b-f) By contrast, spreads of pachytene oocytes containing a mixture of bivalents and univalents expressed satCE02 signals on only two bivalents. This probably corresponded to TT chromosomes (diploid set) and one univalent possibly relating to the E chromosome (haploid set; Figure 4e).

We may therefore conclude that the two types of pachytene cells differ in ploidy. Those with only bivalents emerged from gonocytes with duplicated genomes, which allows all chromosomes to find an identical copy to pair with (Figure 4g right panel; Supplementary Figure 4c right panel, Supplementary Figure 5c right panel). However, those containing a mixture of bivalents and univalents have not passed through genome duplication and their bivalents occasionally formed by homologues *C. taenia* chromosomes from a diploid parental set of chromosomes while orthologous chromosomes of *C. elongatoides* exist as univalent (Figure 4g, left panel; Supplementary Figure 4c left panel, Supplementary Figure 5c left panel).

Additionally, we examined the ploidy of pachytene oocytes in intact gonads of diploid and triploid hybrid females using whole mount FISH with chromosome specific probe satCE01 (Marta et al. 2020). In the somatic cells of diploid hybrids, this marker identifies 1 chromosome of *C. taenia* and 1 chromosome of *C. elongatoides*, while in the somatic cells of triploid ETT hybrids, it determines 2 chromosomes of *C. taenia* and 1 chromosome of *C. elongatoides* (Supplemental Figure 3a,b,e,f) (Marta et al., 2020). This marker therefore allows the distinguishing of the ploidy of pachytenic cells only in triploid hybrids but not diploids not only on chromosomal spreads but also in tissue fragments. Pachytene cells of triploid hybrids with 2 signals would have their genome unduplicated as 1 signal comes from bivalent between 2 TT chromosomes while another signal comes from 1 EE univalent (Figure 5j,k,l). This was exactly the case in cells with a mixture of bivalents and univalents. By contrast, in the case of pachytene cells with bivalents only, we observed 3 signals from 2 TT bivalents and 1 EE bivalent, suggesting their genome was duplicated (Figure 5g,h,i). Such analyses confirmed the aforementioned observations on chromosomal spreads and moreover, we also discovered that pachytene cells with duplicated genomes do not form large clusters but instead occur as individual cells and at most form clusters of 2-4 duplicated cells.

**Fig 5.**
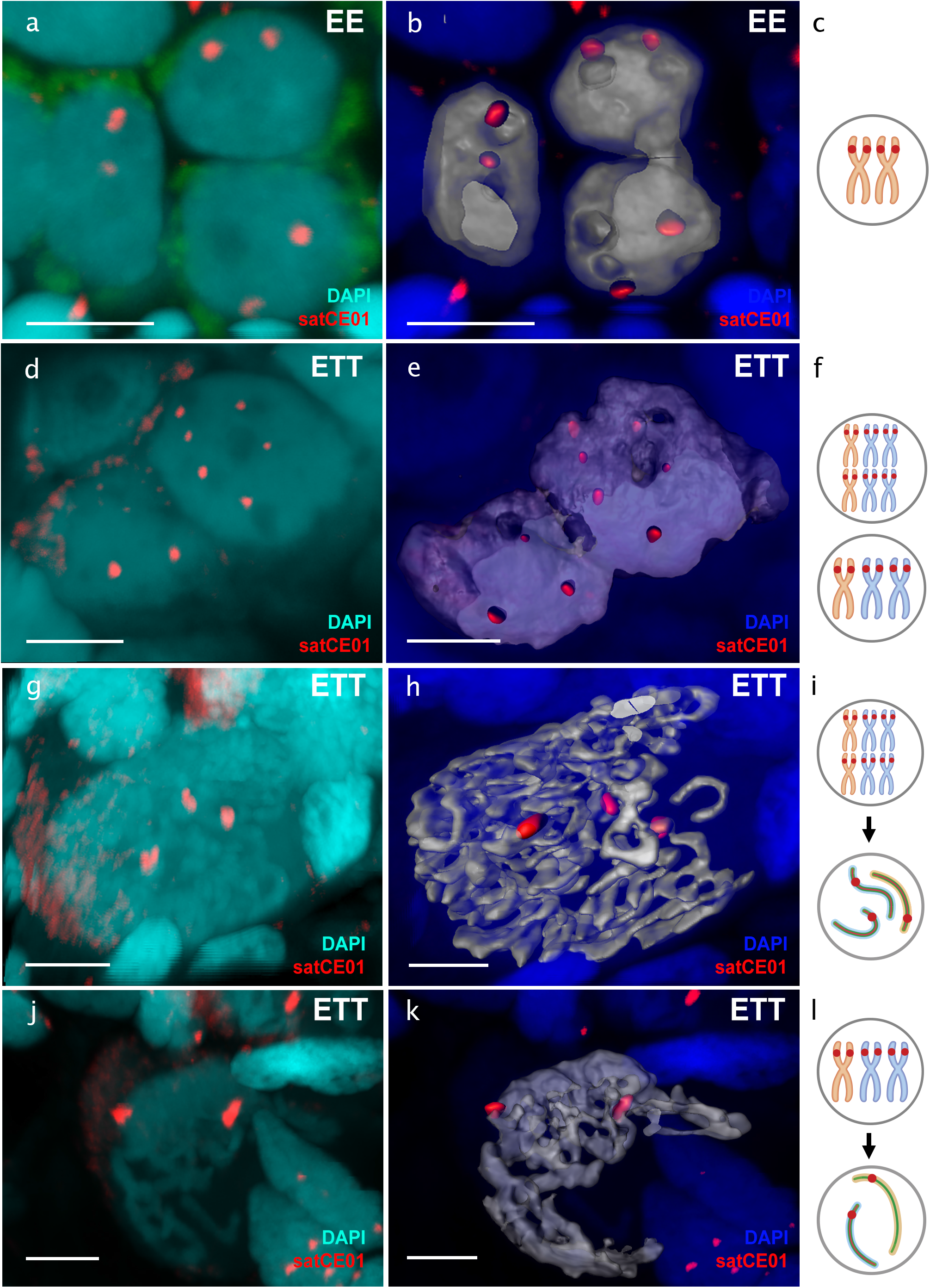
Identification of germ cells ploidy level using FISH with chromosome specific probe in *C. elongatoides*, and triploid hybrids. Whole-mount FISH with chromosome specific probe (a) and corresponding 3D surface reconstruction (b) distinguish 2 chromosomes in germ cells of *C. elongatoides*. Whole-mount FISH with chromosome specific marker (d) and corresponding 3D surface reconstruction (e) identifies 3 chromosomes in germ cells with nonduplicaed genome (triploid cells) and in germ cells with duplicated genome (hexaploid cells) in ovaries from triploid ETT hybrids. (c,f) Schematic representation of karyotype composition of corresponding germ cells. Red marks indicate chromosomes identified by FISH with chromosome specific marker satCE01. Whole-mount FISH with chromosome specific probe (g, j) and corresponding 3D surface reconstructions (h, k) distinguish cells during pachytene with duplicated genome (g, h), and nonduplicated genome (j,k). Images a, c, e, g are 3D reconstructions of natural confocal sections showing the views of germ cells with visualized chromosomes by FISH with chromosome specific marker. Chromosome specific probe stained red, DAPI stained blue. (i) Schematic representation of bivalents and univalents in pachytene cells with duplicated genome (i) and in cells with notduplicated genome (l) as well as presumptive karyotype composition cause such pairing. Red marks indicate bivalents identified by FISH with chromosome specific marker satCE01 as well as presumptive karyotype composition in gonocytes. Scale bar = 5μm.

### Gonocytes (germ cells) also occur in unduplicated and duplicated forms within intact gonad of hybrid females

Finally, we tested whether two ploidy types of cells also exist in the stage of gonocytes. To identify gonocyte cells within the ovarian tissues, we initially applied antibodies against Vasa protein and determined their distinct morphology (Supplementary Figure S6). We subsequently identified the gonocyte’s genome composition using whole mount FISH with chromosome specific marker (satCE01). In sexual species we detected 2 signals per gonial cell (Figure 5a,b,c). In particular, in three triploid fish (ETT) we obtained data on a number of duplicated and nonduplicated cells both at the level of pachytene oocytes as well as gonocytes. We detected on average 89% without genome duplication (cells with 3 signals) and 11% with genome duplication (cells with 6 signals, n=55) (Figure 5d,e,f; Supplementary Table 1). Although the ratio of duplicated cells seemed slightly higher among gonocytes, the difference was not significant (generalized linear mixed effect model with individual taken as a random factor and binomial error distribution, p.value = 0.12).

## Discussion

### Genome duplication is restricted to the minor cell population while the majority of cells may not proceed beyond pachytene

The transition from sexual reproduction to asexuality is often triggered by hybridization and many such hybrid asexual organisms are known to undergo premeiotic endoduplication to produce clonal gametes (Dedukh et al., 2015; Dedukh et al., 2020; Itono et al., 2006; Juchno et al., 2016; Kuroda et al., 2018; Lutes et al., 2010; Macgregor and Uzzell, 1964; Stenberg and Saura, 2009). However, it is largely unknown how widespread this mechanism is within gonial cells of given asexual organisms. To address this, we inspected the genome composition of oocytes during pachytene and diplotene meiotic stages and also of gonial cells in laboratory produced F1 hybrids as well as in natural diploid and triploid asexuals of the *Cobitis* fish. Surprisingly, premeiotic genome endoreplication was observed in only a minor fraction of the hybrid’s gonocytes, while the vast majority of gonial cells were unable to duplicate their genomes causing abruption of pairing and bivalent formation during the pachytene stage (Figure 6). To our knowledge, similar analysis has only been performed in earlier studies, which inferred the rarity of the endoreplication event by examination of the DNA content of gonocytes from parthenogenetic whiptail lizards (from the genus *Aspidoscelis*) and the analysis of meiotic chromosomal spreads in F1 diploid hybrids between two medaka species (Hamaguchi and Sakaizumi, 1992; Newton et al., 2016; Shimizu et al., 2000). Together with our investigation, such data suggest that even successful natural hybrid asexuals suffer from genome incompatibilities and improper chromosomal pairing, while endoreplication, which restores hybrid fertility by allowing chromosomal pairing between identical chromosomal copies, is a rare event in an asexual hybrid’s gametes.

**Fig 6.**
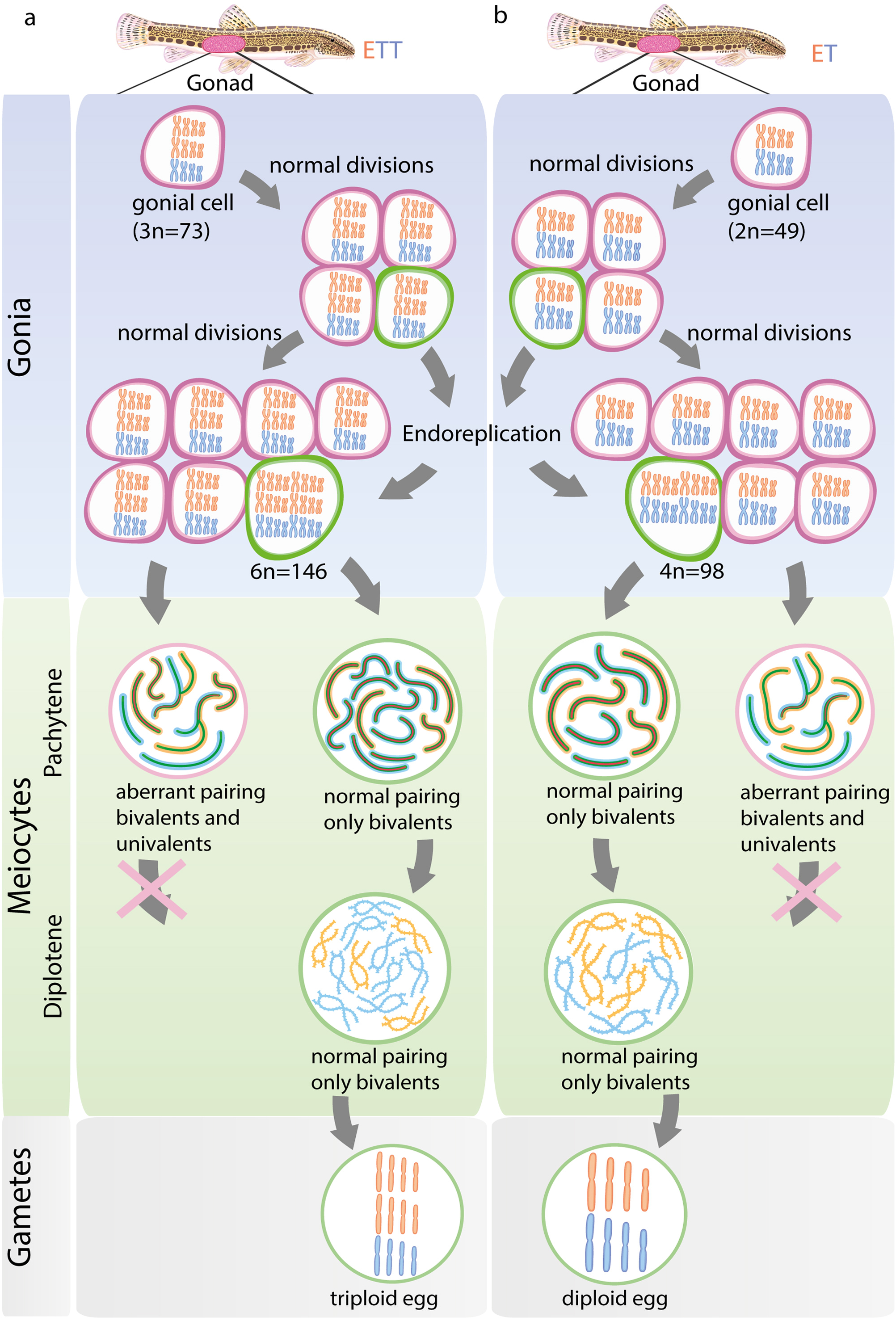
Schematic overview of gametogenic pathway during clonal gametogenesis in triploid ETT (a) and diploid ET (b) hybrids. Each column shows individuals, gametogenic pathways with the indication of the germ cells, cells in two meiotic stages (pachytene and diplotene) and gametes. E (orange chromosomes), T (blue chromosomes) indicates genomes of both parental species *C. elongatoides* and *C. taenia* correspondingly. Cells with premeiotic genome duplication (green outlines) emerged in individuals gonocytes before meiosis. After entering meiosis such cells form 73 bivalents (a) or 49 (bivalents) during pachytene and diplotene followed by the formation of triploid (a) and diploid (b) gametes. Gonocytes with not duplicated genome (grey outlines) enters meiosis and form univalent and bivalent during meiosis. Such cells cannot proceed beyond pachytene.

We further discovered that the ratio of duplicated/nonduplicated cells drastically changed between pachytene and diplotene stages, when oocytes with duplicated genomes and proper bivalents were observed exclusively, without any univalents or misaligned chromosomes (Figure 2). This suggests that oocytes, which may not form proper bivalents without premeiotic endoduplication, cannot proceed beyond the pachytene stage (Figure 6). Such observation coincides with the well-known “pachytene checkpoint” that involves efficient DSB repair machinery and elements controlling synapsis (Roeder and Bailis, 2000; Subramanian and Hochwagen, 2014). As these pathways are highly conserved between yeast, nematodes, insects and mammals (Bohr et al., 2016; Chen et al., 2018; MacQueen and Hochwagen, 2011; Marcet-Ortega et al., 2017), we expect that similar processes take place in *Cobitis* hybrid females and probably in other asexuals too. We may thus propose that DSB-dependent and/or DSB-independent machinery prevent the progression of non-duplicated oocytes with incompletely paired chromosomes beyond pachytene, potentially causing the death of such cells, thereby preventing the growth and formation of aberrant gametes.

### Sex specific differences in meiotic checkpoints and stringency of pairing chromosomal control

Our observations also offer an interesting insight into the sex-specific differences of gametogenic control when compared to previous analysis of hybrid males. Namely, the investigated meiotic divisions in diploid and triploid male hybrids between *C. elongatoides* and *C. taenia*, Dedukh and co-authors (Dedukh et al., 2020) observed aberrant spermatocytes with univalents and multivalents, both in the pachytene stage of meiotic prophase I and in the metaphase of meiosis I. Thus, contrary to hybrid females, male cells with aberrant pairing are fully or partially able to bypass the pachytene checkpoint (Dedukh et al., 2020). Nevertheless, such cells seem to become trapped during the spindle assembly checkpoint acting during metaphase 1 (Lane and Kauppi, 2019; Musacchio and Salmon, 2007). Spindle assembly checkpoint machinery assesses the stringency of the spindle in each bivalent and allows progression beyond metaphase only when all bivalents are correctly arranged (Lane and Kauppi, 2019; Musacchio and Salmon, 2007). Thus, meiotic progression in male hybrids is prevented at later stages by the failure of the equal stringency from the spindle caused by univalents (Burgoyne et al., 2009; Eaker et al., 2002).

Differences between sexes in meiotic checkpoints were also observed in other organisms, but the patterns somewhat contradict each other (Fielder et al., 2020; Kurahashi et al., 2012; Lane and Kauppi, 2019). For example, the hybrids between Medaka fish *Oryzias latipes* × *O. curvinotus* showed similar patterns to *Cobitis* since oocytes with aberrantly paired chromosomes could not proceed beyond pachytene, while spermatocytes with aberrant pairing did not disrupt meiotic prophase but also progressed to metaphase 1 meiosis (Shimizu et al., 2000, 1997). Such abruption of meiosis is probably caused by the abnormal alignment of bivalents at the spindle suggesting that meiotic progression is blocked due to the inability to pass through the spindle assembly checkpoint (Burgoyne et al., 2009; Eaker et al., 2002; Lane and Kauppi, 2019; Musacchio and Salmon, 2007; Shimizu et al., 2000, 1997). By contrast, in asexual triploid hybrids from *Misgurnus*, the sister genus to *Cobitis*, oocytes with univalents and bivalents apparently proceed to diplotene (Zhang et al., 1998). Moreover, they also navigate the spindle assembly checkpoint, as univalents do not attach to the spindle and are lost during the anaphase of meiosis 1 while chromosomes forming bivalents segregate normally (Zhang et al., 1998). Mammals appear yet different from *Cobitis* and other aforementioned cases as the defects in the DSB repair system or synapsis typically lead to cell death during the pachytene stage of mammalian spermatocytes, while in other species, defective oocytes often proceed through both meiotic divisions (Nagaoka et al., 2012, 2011). Taken together, we suggest that having the capacity to overcome different gametogenic pathways (for instance, pachytene or spindle assembly checkpoints) is crucial for the reproduction of asexual hybrid females.

### Initiation of premeiotic genome duplication

A very efficient mechanism to alleviate problems in orthologue pairing and simultaneously gain clonal reproduction is premeiotic genome duplication (this study; Dedukh et al., 2020; Shimizu et al., 2000). Yet, despite its widespread occurrence across major animal and plant taxa, it remains a surprisingly understudied phenomenon with very little known about when the duplication event occurs or how it is triggered.

Our observations can deliver at least some information regarding this. Fish oogonia are able to divide throughout the lifetime of the animal and are maintained as gonial stem cells that can subsequently enter meiosis (Nakamura et al., 2011; Tyler and Sumpter, 1996; Wildner et al., 2013). During their proliferation, the gonocytes actively divide and form clusters (“nests”), that contain all descendants of a single progenitor cell. If endoreplication occurs during the early proliferation stage of gonocytes, large clusters of exclusively endoreplicated cells would be expected. However, our analysis of pachytene cells found no evidence of such clusters as we mostly observed individual nuclei with duplicated chromosomal sets, or rarely small groups of 2 - 4 endoreplicated cells (Figure 4c,d; Figure 5d,e). We may thus conclude that endoreplication occurs in gonocytes just before entering meiosis or occasionally 1-2 divisions before such an event.

In *Misgurnus* loaches, Yoshikawa and co-authors (Yoshikawa et al., 2007) described a different timing of endoreplication in the gonocytes of hybrid males hormonally reversed into females. They found that endoreplication occurred in sex-reverted hybrid males already in A-type spermatogonia, which somewhat contradicts our observation in *Cobitis* females, since such spermatogonia still have several mitotic divisions before turning into B-type spermatogonia and thus, entering meiosis. Such a comparison suggests that premeiotic endoreplication may proceed differently even in closely related organisms such as *Cobitis* and *Misgurnus*. Alternatively, the initiation of endoreplication may be caused by methodological approaches and these contrasting results could be obtained for instance by the developmental shock associated with sex-reversal as suggested by Yoshikawa et al. (Yoshikawa et al., 2007).

While detecting the timing of endoreplicaton is challenging, it is even harder to identify the causal triggers initiating this process in asexual hybrids. Interestingly however, three independent studies on unrelated organisms documented that endoreplication affects only a rather minor fraction of gonial cells in asexual hybrids (such as lizards and at least two fish species [this study; Hamaguchi and Sakaizumi, 1992; Newton et al., 2016; Shimizu et al., 2000]). This may indicate that some similarities exist in underlying mechanisms causing this aberration. We therefore believe that understanding of the mechanisms causing genome duplication in germ cells of asexual hybrids can be improved based on studies implemented on polyploid/aneuploid cells observed in a variety of organisms e.g. flies, mice, humans and the zebrafish (Calvi, 2013; Fox and Duronio, 2013; Lee et al., 2009; Orr-Weaver, 2015). For instance, it has been documented that endopolyploidy affecting some cells or tissues emerge during development or under stress conditions (Cao et al., 2017; González-Rosa et al., 2018; Losick et al., 2013). In different organisms, several cell cycle regulators were found to be responsible for the switch from normal mitosis to endomitosis/ endoreplication cycles, specifically when they are downregulated or their function is affected (Diril et al., 2012; Nannas and Murray, 2012; Rotelli et al., 2019; Sauer et al., 1995). Among these regulators, Cyclin A/Cyclin dependent kinase 1 and Aurora B kinase play a crucial role (Adams et al., 2001; Diril et al., 2012; Giet and Glover, 2001; Nannas and Murray, 2012; Rotelli et al., 2019; Sauer et al., 1995). Moreover, in their elegant work, Rotelli and co-authors (Rotelli et al. 2019) showed the link between concentration of CDC1 and AurB to the stringency of effects on the cell cycle. Strong knockdown of CDK 1 or AurB in human cancer cells and *Drosophyla* embryos inhibited chromosome segregation and cytokinesis causing G-S cycles (endoreplication), whereas a mild knockdown resulted in successful chromosome segregation but failure of cytokinesis (endomitosis) (Chen et al., 2016; Rotelli et al., 2019). We may therefore hypothesize that in hybrids some proteins involved in the Cyclin/Cdk1—AurB pathway can be translated from genomes of different parental species forming enzyme heterocomplexes with a decreased function as compared proteins composed of products from the same species. Such heterocomplexes may then lead to a decreased level of AurB kinase causing a round of endoreplication/endomitosis in individual gonocytes of hybrid females.

### Implications for ecology and evolution of asexual organisms

The emergence of gametogenic aberrations leading to asexuality was suggested to be either directly triggered by hybridization (Schultz, 1969) or as a result of preconditions in sexual species with hybridization just accelerating this process (Cuellar, 1974; Sinclair et al., 2010). One way or another, asexual reproduction provides a considerable short-term advantage by avoiding the cost of male offspring and, all else being equal, asexuals should quickly outcompete their sexual counterparts. In the reality, however, sexual and asexual counterparts differ by many traits. For example, asexuality is often linked to polyploidy, which modifies the metabolic rate (Maciak et al., 2011), or size-selective predation mortality (Juchno et al., 2014, 2013) when a larger size of polyploid offspring allows faster escape from natural predators (Sogard, 1997). In any case, the extinction of sexuals would leave sperm-dependent asexuals like gynogens without a source of sperm and hence some mechanisms are clearly necessary in order to stabilize sexual-asexual coexistence. These usually assume some behavioural adaptations linked to mate choice (Mee and Otto, 2010; Morgado-Santos et al., 2015; Schlupp and Plath, 2005), but it has been suggested that increased mortality or sterility of asexual individuals may be beneficial for gynogenetic populations since it would compensate the inherent demographic advantage of asexuality and stabilize their coexistence with sexual species (Bobyrev et al., 2003; Leung and Angers, 2018). In the *Cobitis* hybrid complex, Bobyrev et al. (Bobyrev et al., 2003) and Juchno and Boroń (Juchno and Boroń, 2010, 2006) reported an approximately 50% reduction in fecundity in triploid *elongatoides-taenia* hybrid females compared to the *C. taenia* parental species indicating the potential existence of such compensation in natural populations. On the other hand, the reported drop in fecundity is far lower than our observation of failure in pachytene oocytes, suggesting that some cellular mechanism(s) compensate the dramatic loss at the pachytene checkpoint.

Although possibly advantageous at population level, such a high rate of gametogenic failure is disadvantageous for individuals and it is likely that short-term selection should favour those hybrid clones that minimize the negative effect of genomic incompatibilities. Our data support such a hypothesis as we found a significantly higher rate of endoreduplication in successfully established natural clones as compared to experimental F1 clones originated from crossings of randomly selected parents. Thus, although initiation of asexuality is achieved frequently by hybridization of *C. elongatoides* with other species (this study; Choleva et al., 2012), interclonal selection appears to favour hybrid clones which originated from particular combinations of parental genomes allowing the highest rates of endoreduplication. It is also possible that after their formation some clones evolve to further ameliorate such capability. On the other hand, although the most widespread and oldest *Cobitis* clone (EEN) has a significantly higher fecundity compared to relatively younger ET and ETT lineages that originated in Central Europe during the Holocene. Earlier, we found no evidence for any higher rate of endoreduplication in this successful lineage, suggesting that selection may possibly also operate on later stages of gametogenic processes (Kočí et al., 2020).

## Conclusion

In conclusion, asexuality and hybrid sterility are intimately intertwined phenomena and transitions from sexual reproduction to asexuality may bring considerable costs, especially in hybrid taxa which face significant problems with genome incompatibilities, potentially greatly reducing their reproductivity (Figure 6). Nevertheless, asexuals may establish very successful and persistent lineages even with such an expense, possibly helping them to maintain a stable coexistence with sexual hosts. Our study also stresses the importance of using several approaches experimentally, as demonstrated by the striking differences in cell ploidy detected in pachytene and diplotene stages that were detected by different experimental means. It should be noted that taking into account only one type of analysis could lead to misinterpretation of the results. Finally, it also appears of crucial importance to collect detailed insights into gametogenic pathways of various asexuals in order to understand the mechanistic interlink between hybridisation, sterility and asexuality.

## M&M

### Samples studied

Genome composition and ploidy of every investigated specimen were evaluated with the set of species-diagnostic markers including microsatellites, allozymes, and cytogenetic methods (Janko et al. 2007). In total, we analysed 9 triploid ETT hybrid females, 4 triploid EEN hybrid females and 11 diploid hybrid females from nature localities and laboratory crosses. Two individuals of parental species (*C. elongatoides*) were used as a control. Any treatment or injection was used before the investigation of female gametogenesis. Animals were anesthetized in MS222 and sacrificed accordingly. Gonads of each individual were separated into several pieces followed by usage for pachytene and/or diplotene chromosome preparation and for observation under laser scanning confocal microscope. Gonadal tissues used for 3D analysis were fixed in 2% paraformaldehyde in 1× PBS for 90 min at room temperature (RT), washed in 1× PBS, and kept in 1 × PBS with 0.02% NaN3 until usage.

### Crossing experiments

Spawning was performed artificially by hormonal injections of males and females. Injection of hormone Ovopel (Interfish Kf) was performed twice in the peritoneal cavity. First injection (24 hours before fertilization) was applied with a solution of one Ovopel pill per 20 mL of 0.9% NaCl. Second injection (12 hours before fertilization) was done with a solution of 1 pill per 5 mL of 0.9% NaCl. The ratio of injected solution in both cases was 0.05 mL per 10g of fish weight. Fish eggs were gently squeezed 24 hours after first injection and transferred to a Petri dish. Male sperm was also obtained by squeezing and directly applied on loach eggs with the addition of fresh water to activate spermatozoa. After hatching, free larvae were transferred into plastic pots (25 × 25 × 15 cm). After two-three month after hatching we randomly selected juveniles for the analysis of pachytene oocytes. Diplotene oocytes were analysed from adult and subadult females older half of the year.

### Pachytene chromosomes and immunofluorescent staining

Pachytene chromosomes were obtained according to protocols described by Araya-Jaime et al.55. After manual homogenization of female gonads, 20 μl of cells suspension was dropped on SuperFrost^®^ slides (Menzel Gläser) followed by addition of 40 μl of 0.2 M Sucrose and 40 μl of 0.2% Triton X100 for 7 min. Afterward, cells were fixed for 16 minutes by adding 400 μl of 2% PFA. After washing in 1 × PBS slides were stored until immunofluorescent staining of synaptonemal complexes (SC) was performed.

Lateral components of synaptonemal complexes (SC) were detected by rabbit polyclonal antibodies (ab14206, Abcam) against SYCP3 protein while the central component of SC was detected by chicken polyclonal SYCP1 (gift from Prof. Sean Burgess). According to previously published data SYCP3 is localized on both on bivalents and univalents while SYCP1 indicates pairing and bivalent formation (Blokhina et al 2019). Fresh slides were incubated with 1% blocking reagent (Roche) in 1 × PBS and 0.01% Tween-20 for 20 min followed by the addition of primary antibody for 1h at RT. Slides were washed 3 times in 1× PBS at RT and incubated in a combination of secondary antibodies (Cy3-conjugated goat anti-rabbit IgG (H+L) (Molecular Probes) and Alexa-488-conjugated goat anti-mouse IgG (H+L) (Molecular Probes) diluted in 1% blocking reagent (Roche) on PBS for 1h at RT. Slides were washed in 1× PBS and mounted in Vectashield/DAPI (1.5 mg/ml) (Vector, Burlingame, Calif., USA).

### Diplotene chromosomes

Diplotene chromosomal spreads (so-called “lampbrush chromosomes”) were prepared from parental and hybrid females according to an earlier published protocol (XXX). Ovaries from non-stimulated females were dissected and placed in the OR2 saline (82.5 mM NaCl, 2.5 mM KCl, 1 mM MgCl2, 1 mM CaCl2,1mM Na2HPO4, 5 mM HEPES (4-(2-hydroxyethyl)-1-piperazineethanesulfonic acid); pH 7.4). Oocyte nuclei were isolated manually using jeweller forceps (Dumont) in the isolation medium “5:1” (83 mM KCl, 17 mM NaCl, 6.5 mM Na2HPO4, 3.5 mM KH2PO4, 1mM MgCl2, 1 mM DTT (dithiothreitol); pH 7.0–7.2). Oocyte nuclei were transferred to glass chambers attached to a slide filled in one-fourth strength “5:1” medium with the addition of 0.1% paraformaldehyde and 0.01% 1M MgCl2. Such a method ensures that each chamber contained chromosomal spread from the individual oocyte. The slide was subsequently centrifuged for 20 min at +4°C, 4000 rpm, fixed for 30 min in 2% paraformaldehyde in 1× PBS, and post-fixed in 70% ethanol overnight (at +4°C).

### Fluorescence *in situ* hybridization

Probes for fluorescent in situ hybridization (FISH) procedures were selected according to earlier published data (Marta et al., 2020). We selected chromosome-specific markers: (satCE01), centromeric repeat (satCE04); as well as a species polymorphic marker (satCE02) between *C. elongatoides* and *C. taenia*. Biotin and digoxigenin labelling of probes were performed by PCR using genomic DNA of *C. elongatoides* according to Marta et al., 2020.

The hybridization mixture contained 50% formamide, 10% dextran sulfate, 2× SSC, 5 ng/μl labeled probe, and 10–50-fold excess of tRNA. After common denaturation of the probe and chromosomal DNA on slides at 75°C for five minutes, slides were incubated for 12-24 hours at room temperature (RT). After hybridization, slides were 3 times washed in 0.2× SSC at + 44°C. The biotin-dUTP and digoxigenin-dUTP were detected using streptavidin-Alexa 488 (Invitrogen, San Diego, Calif., USA) and anti-digoxigenin-rhodopsin (Invitrogen, San Diego, Calif., USA) correspondingly. The chromosomes were counterstained with Vectashield/DAPI (1.5 mg/ml) (Vector, Burlingame, Calif., USA).

### Whole-mount immunofluorescence staining

Prior immunofluorescent staining, gonadal fragments were permeabilized in a 0.5% solution of Triton X100 in 1× PBS for 4-5 hours at RT followed by washing in 1× PBS at RT. Following primary antibodies were used rabbit polyclonal antibodies DDX4 antibody [C1C3] (GeneTex) against Vasa protein; rabbit polyclonal antibodies against SYCP3 (ab14206, Abcam); chicken polyclonal against SYCP1 (gift from Prof. Sean Burgess; Blokhina et al 2019). After incubation for 1-2 hours in a 1% blocking solution (Roche) dissolved in 1× PBS, primary antibodies were added for 12 hours at RT. Secondary antibodies anti-rabbit conjugated with Alexa-488 fluorochrome and anti-chicken conjugated with Alexa 594 for 12 hours at RT were used. Washings from primary and secondary antibodies were carried out in 1× PBS with 0.01% Tween (ICN Biomedical Inc). Tissues were stained with DAPI (1 μg/μl) (Sigma) in 1× PBS at RT overnight.

### Whole-mount fluorescence *in situ* hybridization

Gonadal fragments were permeabilized in 0.5% solution of Triton X100 in 1× PBS for 4-5 hours at RT followed by impregnation by 50% formamide, 10% dextran sulfate, and 2×SSC for 3-4 hours at 37°C. Afterward, tissues were placed to hybridization mixture including 50% formamide, 2× SSC, and 10% dextran sulfate, 20 ng/μl probe, and 10 to 50-fold excess of salmon sperm DNA. Gonadal tissues were denaturated at 82°C for 15 minutes and incubated for 24 hours at RT. Tissues were washed in three changes of 0.2× SSC at 44°C for 15 minutes each and blocked in 4×SSC containing 1% blocking reagent (Roche) in 4× SSC for 1 hour at RT. The biotin-dUTP and digoxigenin-dUTP were detected using streptavidin-Alexa 488 (Invitrogen, San Diego, Calif., USA) and anti-digoxigenin-rhodopsin (Invitrogen, San Diego, Calif., USA) correspondingly. The tissues were stained with DAPI (1 mg/ml) (Sigma) diluted in 1× PBS at RT overnight.

### Confocal laser scanning microscopy

Tissues fragments were placed in a drop of DABCO antifade solution containing 1 mg/ml DAPI. Confocal laser scanning microscopy was carried out using a Leica TCS SP5 microscope based on the inverted microscope Leica DMI 6000 CS (Leica Microsystems, Germany). Specimens were analysed using HC PL APO 40x objective. Diode, argon and helium-neon lasers were used to excite the fluorescent dyes DAPI, fluorochromes Alexa488, and Cy3, respectively. The images were captured and processed using LAS AF software (Leica Microsystems, Germany).

### Wide-field and fluorescence microscopy

Meiotic chromosomes after FISH and IF were analysed using Provis AX70 Olympus microscopes equipped with standard fluorescence filter sets. Microphotographs of chromosomes were captured by CCD camera (DP30W Olympus) using Olympus Acquisition Software. Microphotographs were finally adjusted and arranged in Adobe Photoshop, CS6 software; Corel Draw GS2019 was used for scheme drawing. Imaris 7.7.1 (Bitplane) software was used for the 3D-volume and surface reconstruction of confocal image stacks. The image stacks used for reconstruction were cropped to the region of interest and used for the reconstruction of isosurfaces. Following channels were used to construct isosurfaces separately: DAPI channel and rhodopsin channel. Threshold parameters were selected automatically. To highlight the visualization of germ cells, only surface objects belonging to individual germ cells were retained in the reconstruction.

## Acknowledgements

Authors would like to thank Dr. Joerg Bohlen (Institute of Animal Physiology and Genetics, Czech Academy of Science) for the significant help with fish care; prof. Sean Burgess (College of Biological Sciences, University of California) for the providing primary antibodies against SYCP1 and critical comments of the manuscript; Antonina Maslova (Saint Petersburg State University) for the help with 3D imaging and Vladislav Vasiulin for the help with the preparation of illustrations.

## Authors’ contributions

DD, AM and KJ conceived the study and designed the experiments. AM and DD performed crosses experiments and preparation of tissue collection. AM performed species identification. DD and AM carried out cytogenetic experiments, whole-mount *in situ* hybridization and whole mount immunofluorescence staining. DD obtained and analyzed confocal images. DD have written the first draft of the manuscript which was further improved by KJ and AM.

## Data availability

The authors state that all data necessary for confirming the conclusions presented in the article are represented fully within the article.

