## Supplementary figures S1-S6; Supplementary table S1 for "Challenges and costs of asexuality: Variation in premeiotic genome duplication in gynogenetic hybrids from *Cobitis taenia* complex": Supplementary files.pdf

**Fig S1. FISH based identification of bivalents from diplotene oocytes of diploid ET female.** (a) Full diplotene chromosomal spread including 49 bivalents. Since the chromosomal spread from individual oocyte was large, eight images were taken and merged into one. Asterisks indicate enlarged bivalents represented on b-d panel. (b-d) S cale bar = 50 µm.

(b-d) High-resolution mapping of species olymor ic at red and centromeric at reen repeats on chromosomes from diplotene oocyte (so-called "lampbrush chromosomes") indicates one bivalents of *C. elongatoides* (b) and 2 bivalents of *C. taenia* (c,d). Scale bar = 5 µm.

(e) Schematic representation of gametogenic pathway, which results in the formation of bivalents observed during diplotene. Purple marks indicate bivalents identified by FISH with species polymorphic marker atCE02 as well as presumptive karyotype composition in gonocytes.

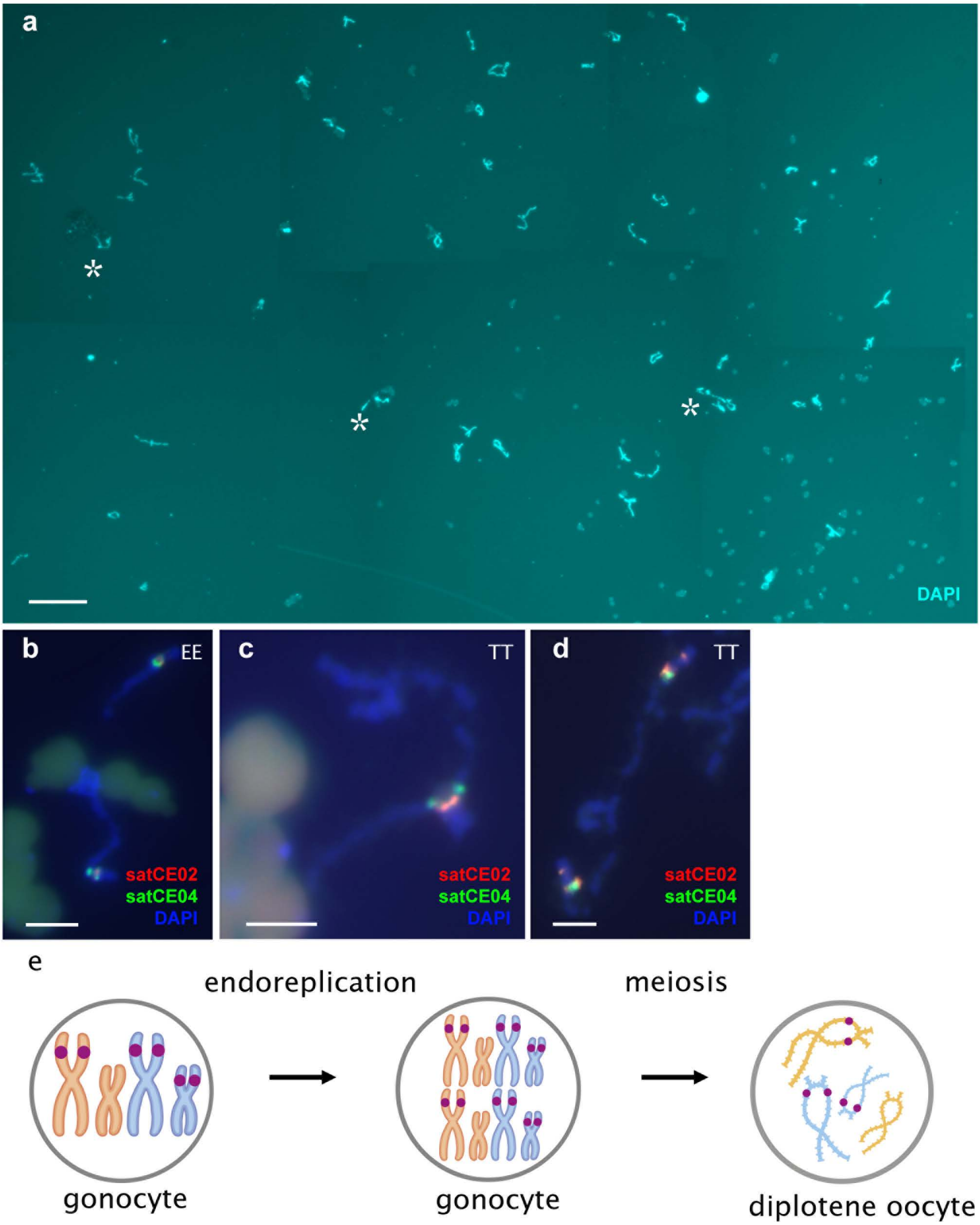
a
DAPI
b
EE
satCE02
satCE04
DAPI
c
TT
satCE02
satCE04
DAPI
d
TT
satCE02
satCE04
DAPI
e
endoreplication
meiosis
gonocyte
gonocyte
diplotene oocyte

**Fig S2. Chromosomal spread from diplotene oocyte of EEN female.** (a) Full diplotene chromosomal spread including 75 bivalents. Since the chromosomal spread from individual oocyte was large, 12 images were taken and merged into one. Scale bar = 50μm. (b) Schematic representation of gametogenic pathway, which results in the formation of bivalents observed during diplotene.

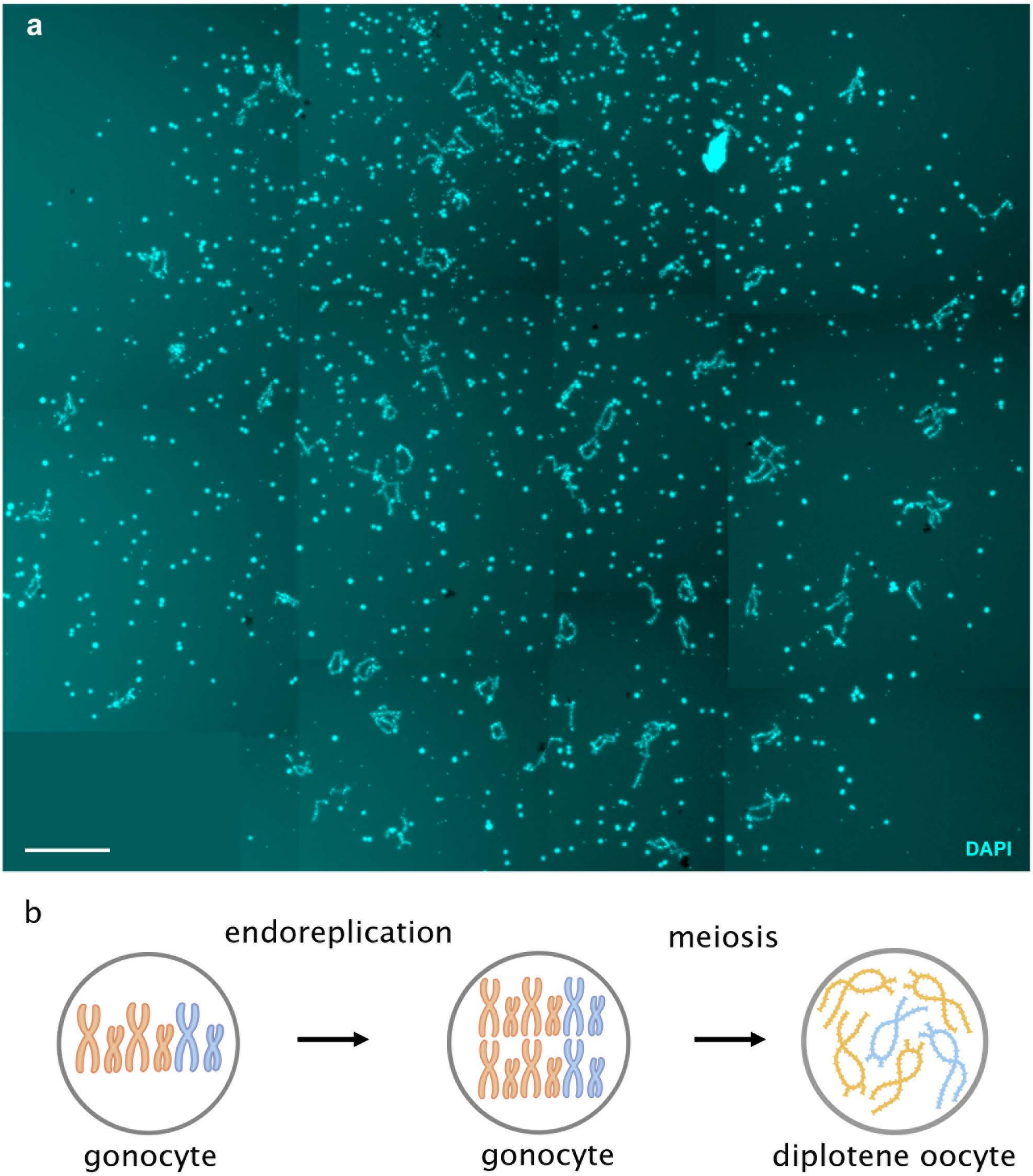
a
DAPI
b
endoreplication
meiosis
gonocyte
gonocyte
diplotene oocyte

**Fig S3 Mapping of satDNA markers on chromosomes of *C. taenia* and *C. elongatoides*.** (a,e) chromosome specific marker SatCE01, and (c,g) species polymorphic marker SatCE02. (b,d,f,h) schematic representation of chromosomes distinguished by satDNA markers within karyotype of *C. elongatoides* (f, h, marked orange) and *C. taenia* (b,d, marked blue). Red marks indicates chromosome specific marker (SatCE01); purple marks indicates species polymorphic marker (Sat CE02).

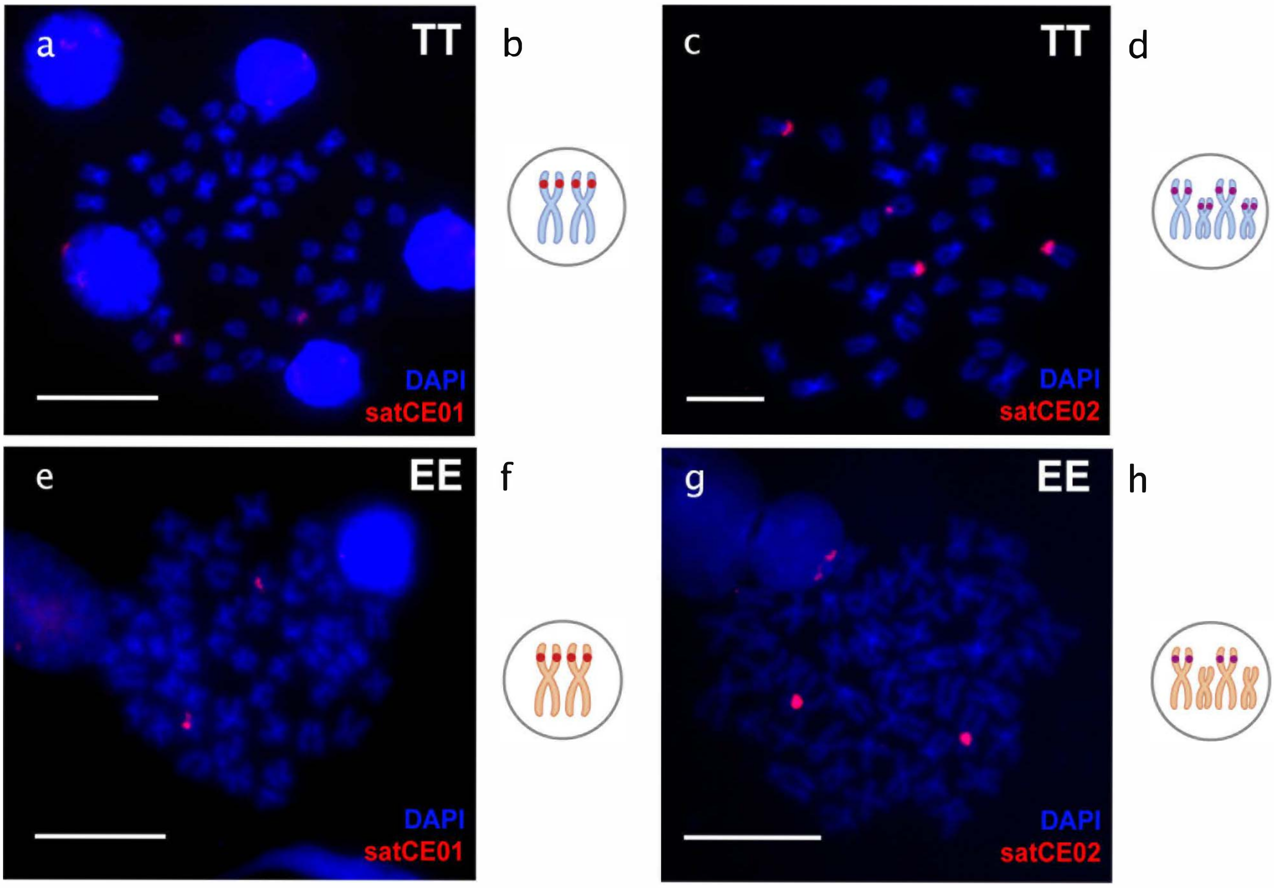
a
TT
DAPI
satCE01
b
c
TT
DAPI
satCE02
d
e
EE
DAPI
satCE01
f
g
EE
DAPI
satCE02
h

**Fig S4. Meiotic cells at pachytene stage from ovaries of diploid ET females.** Visualization of synaptonemal complexes using immunolabeling with antibodies against SYCP3 protein (green) and SYCP1 (red) stained with DAPI (blue). ivalent o ot and locali ation ile univalent e i it only tainin ac ytene oocyte it not du licated enome a and du licated enome Schematic representation of gametogenic pathways in two populations of pachytene cells c . Scale bar = 10µm.

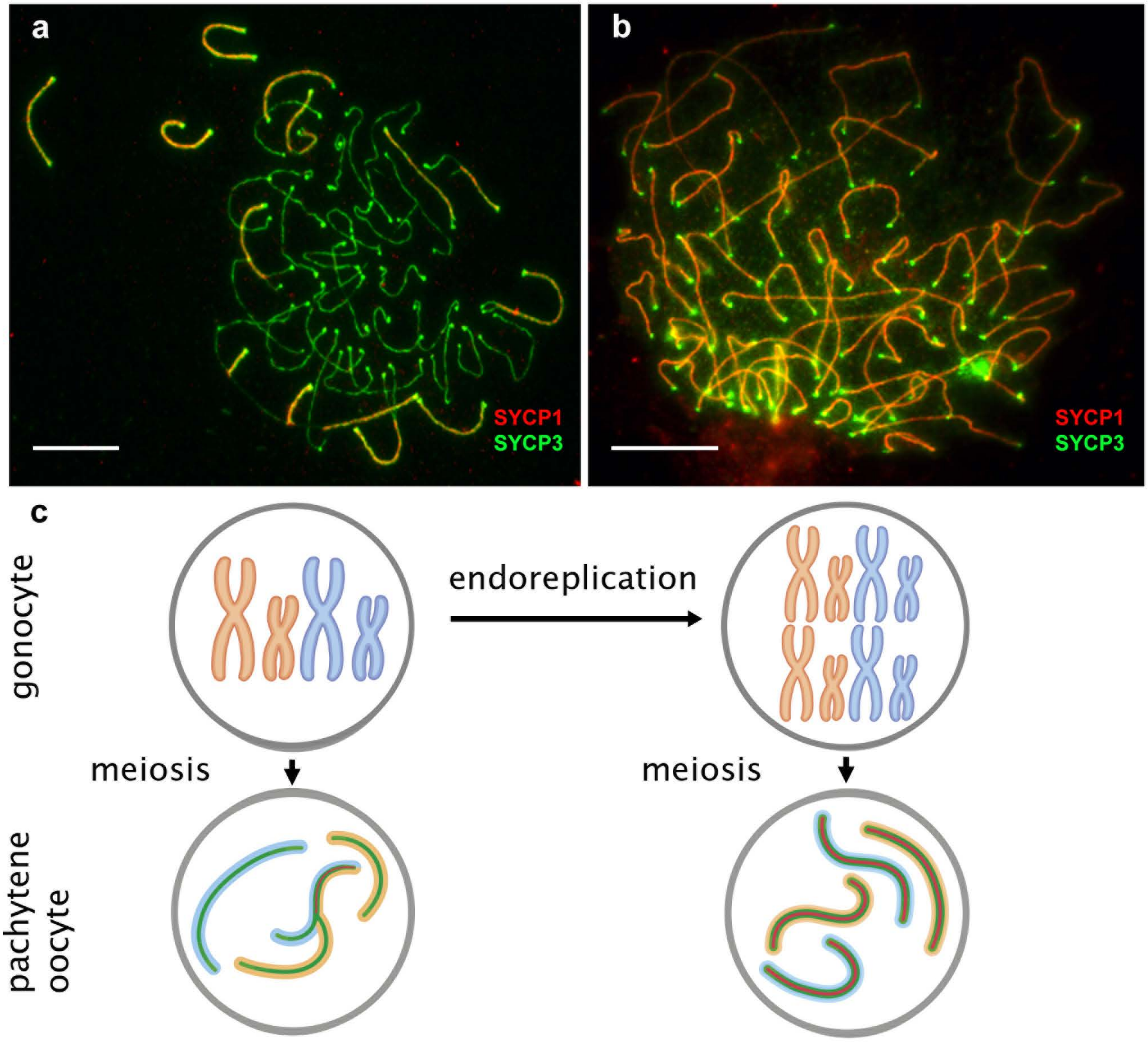
a
SYCP1
SYCP3
b
SYCP1
SYCP3
c
gonocyte
endoreplication
meiosis
meiosis
pachytene oocyte

**Fig S5 Meiotic cells at pachytene stage from ovaries of triploid EEN females.** (a, b) Visualization of synaptonemal complexes using immunolabeling with antibodies against SYCP3 protein (green) and SYCP1 (red) stained with DAPI (blue). Bivalents show both SYCP3 and SYCP1 localization while univalents exhibit only SYCP3 staining.   ac  ytene oocyte     it   not du  licated   enome   a   and du  licated   enome         c  ematic re  re  entation of   ameto  enic   at     ay   in t   o   o  ulation   of   ac  ytene cell    c   Scale bar = 10μm.

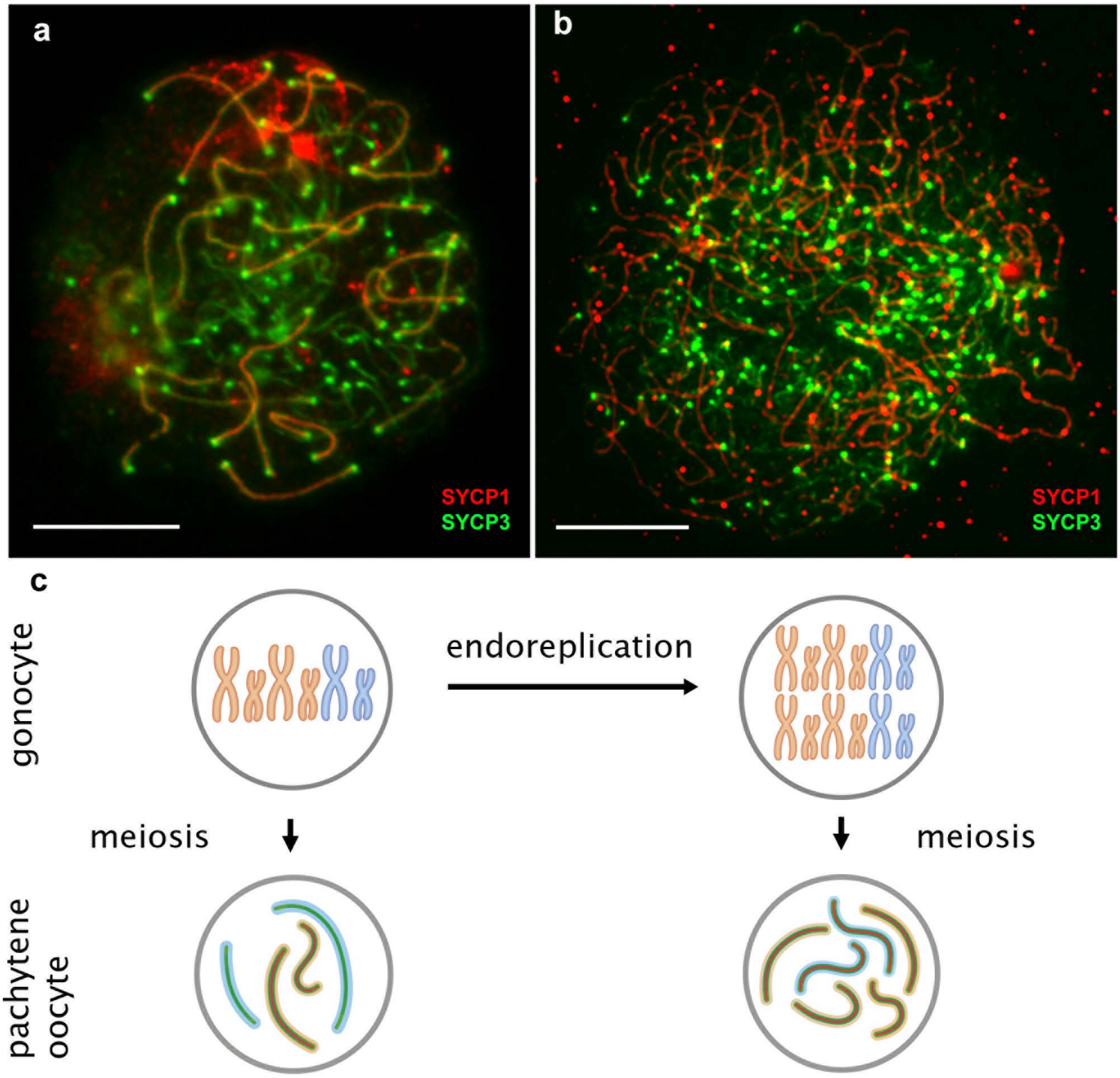
a
SYCP1
SYCP3
b
SYCP1
SYCP3
c
gonocyte
endoreplication
meiosis
meiosis
pachytene
oocyte

**Fig S6 Germ cells in the gonads of triploid ETT hybrids identified using anti-vasa (red) antibodies.** cale ar m

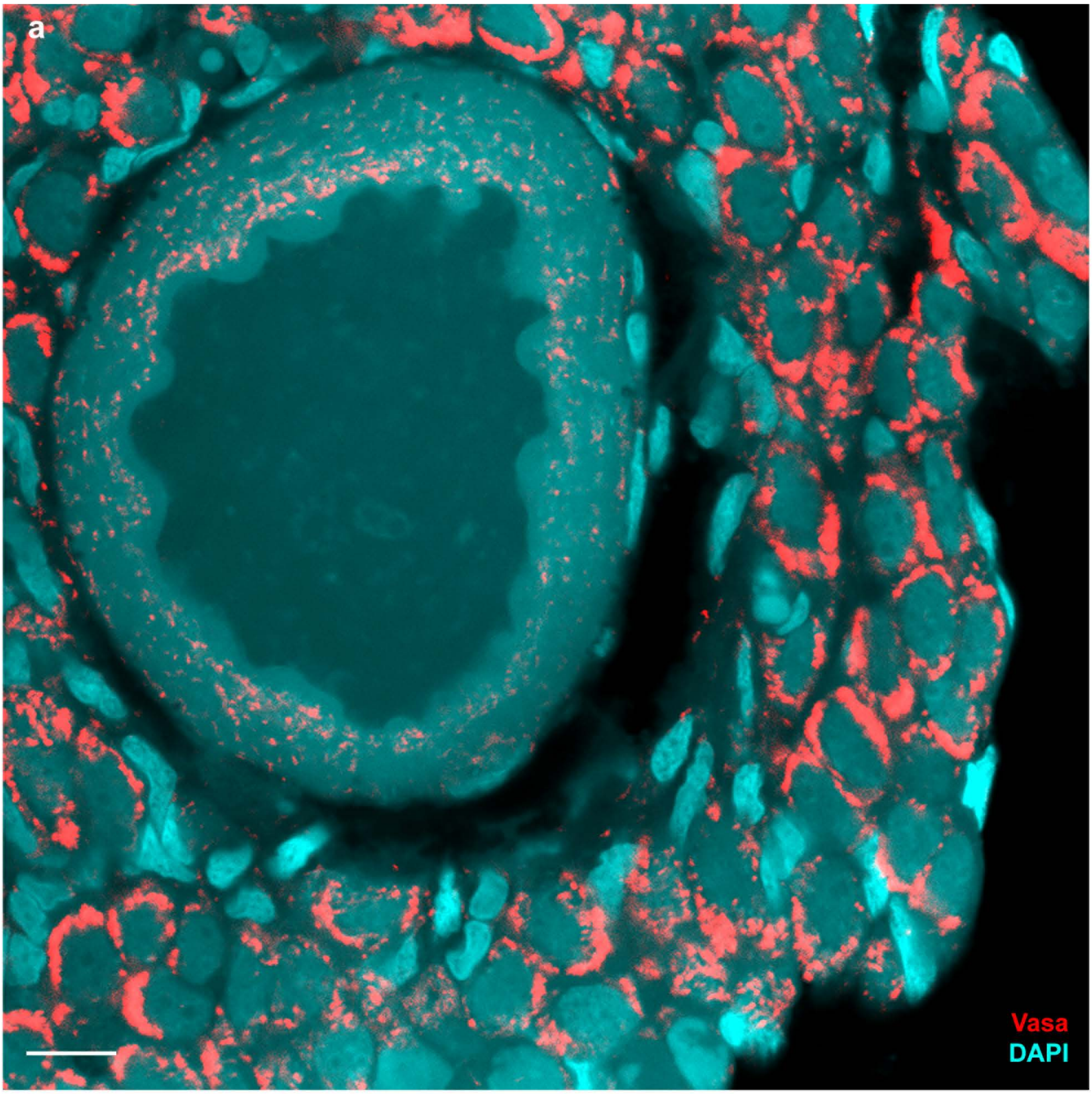
a
Vasa
DAPI

**Supplementary Table S1. List of hybrid individuals from *Cobitis* genus used in the study with detailed description of duplicated and not duplicated cells.**

| | | | pachytene spreads | | diplotene spreads | | IF pachytenes confocal | | FISH pachytenes confocal | | FISH germ cells confocal | |
|---|---|---|---|---|---|---|---|---|---|---|---|---|
| # | Individuals | Genotype | Number of cells with not duplicated genome | Number of cells with duplicated genome | Number of cells with not duplicated genome | Number of cells with duplicated genome | Number of cells with not duplicated genome | Number of cells with duplicated genome | Number of cells with not duplicated genome | Number of cells with duplicated genome | Number of cells with not duplicated genome | Number of cells with duplicated genome |
| | | | pachytene spreads | | diplotene spreads | | IF pachytenes confocal | | FISH pachytenes confocal | | FISH germ cells confocal | |
| 1 | ETT 1 | ETT | 15 | 0 | N/A | N/A | | | | | | |
| 2 | ETT 2 | ETT | 32 | 1 | N/A | N/A | | | | | | |
| 3 | ETT 3 | ETT | N/A | N/A | 0 | 17 | | | | | | |
| 4 | ETT 4 | ETT | 17 | 2 | 0 | 15 | | | | | | |
| 5 | ETT 5 | ETT | 24 | 0 | 0 | 31 | | | | | | |
| 6 | ETT 6 | ETT | 31 | 2 | N/A | N/A | N/A | N/A | 40 | 4 | 152 | 8 |
| 7 | ETT 7 | ETT | 19 | 0 | N/A | N/A | 62 | 6 | N/A | N/A | N/A | N/A |
| 8 | ETT 8 | ETT | 7 | 0 | N/A | N/A | 18 | 4 | 32 | 3 | 237 | 25 |
| 9 | ETT 9 | ETT | N/A | N/A | no | 32 | N/A | N/A | 26 | 4 | 95 | 22 |
| 1 | ET 1 | ET | 11 | 0 | 0 | 27 | | | | | | |
| 2 | ET 2 | ET | 44 | 2 | 0 | 35 | | | | | | |
| 3 | ET 3 | ET | N/A | N/A | 0 | 18 | | | | | | |
| 4 | ET 4 | ET | N/A | N/A | 0 | 18 | | | | | 7 | 1 |
| 5 | ET 5 | ET | 33 | 11 | N/A | N/A | | | | | | |
| 6 | ET 6 | ET | 19 | 1 | N/A | N/A | | | | | | |
| 7 | ET 7 | ET | 46 | 7 | N/A | N/A | | | | | | |
| 8 | ET 8 | ET | 154 | 0 | N/A | N/A | | | | | | |
| 9 | ET 9 | ET | 11 | 1 | N/A | N/A | | | | | | |
| 10 | ET 10 | ET | 53 | 2 | N/A | N/A | | | | | | |
| 11 | ET 11 | ET | 9 | 0 | N/A | N/A | | | | | | |
| 1 | F1_ET 1 | F1_ET | 51 | 0 | N/A | N/A | | | | | | |
| 2 | F1_ET 2 | F1_ET | 43 | 0 | N/A | N/A | | | | | | |
| 3 | F1_ET 3 | F1_ET | 108 | 1 | N/A | N/A | | | | | | |
| 4 | F1_ET 4 | F1_ET | 21 | 0 | N/A | N/A | | | | | | |
| 5 | F1_ET 5 | F1_ET | N/A | N/A | 0 | 15 | | | | | | |

| | | | | | | |
|---|---|---|---|---|---|---|
| 6 | F1_ET 6 | F1_TE | 3 | 0 | N/A | N/A |
| 7 | F1_ET 7 | F1_TE | 72 | 0 | N/A | N/A |
| 8 | F1_ET 8 | F1_TE | N/A | N/A | 0 | 9 |

| | | | | | | |
|---|---|---|---|---|---|---|
| 1 | EEN1 | EEN | N/A | N/A | 0 | 31 |
| 2 | EEN2 | EEN | N/A | N/A | 0 | 15 |
| 3 | EEN3 | EEN | 31 | 1 | N/A | N/A |
| 4 | EEN4 | EEN | 8 | 0 | N/A | N/A |
